# Decoding Ten Years of Little Penguin Foraging: Bio-Logging Reveals Foraging Patterns with Implications for Climate Change Mitigation and Marine Spatial Planning

**DOI:** 10.1101/2023.10.14.562344

**Authors:** Marianna Chimienti, Andre Chiaradia, Benjamin Dupuis, Nicolas Joly, Claire Saraux, Yan Ropert-Coudert, Akiko Kato

## Abstract

1. Protected areas are a widely adopted resource management strategy for mitigating the consequences of global change and preserve functioning ecosystems. Long-term species monitoring programmes, aided by bio-logging technology, provide insights into the extent and spatial variation of areas occupied by wild animals and inform conservation and management. High-resolution GPS-acceleration data offer a more accurate understanding of animal behavior and area use, compared to location-based inference, emphasizing the significance of specific sites amid long-term climate change.
2. We based our case-study on the largest colony of little penguins (*Eudyptula minor*) located at Phillip Island. Based on a ten-year bio-logging dataset (247 individual tracks), we combine high-resolution bio-logging data from GPS-accelerometer loggers with proxies for resource availability (e.g. Sea Surface Tenperature, thermocline, water turbidity). Using machine learning techniques and Generalized Additive Mixed Models, we quantify the environmental factors determining spatio-temporal variability in foraging effort (defined as hunting time) across different breeding seasons and stages.
3. Little penguins increased their hunting time by reducing spatial displacement (shorter step length) and diving deeper, with a slower increase in hunting effort below 10 m depth. In relation to environmental conditions, penguins increased hunting effort in coastal areas with high turbid and productive waters and decreased effort with increasing Sea Surface Temperature. This gives insights into how these animals allocate effort differently according to shifting environmental conditions.
4. Our analysis offers crucial long-term insights into little penguin area usage in the Bass Strait at sufficient spatial and temporal resolution for management and conservation planning. The Bass Strait is facing intense climatic and anthripogenic pressures, and the findings here on intensity of area usage and strategy shifting according to environmental conditions, are of great relevance for the marine spatial planning currently under development along the coast.
5. *Policy implications:* High-resolution behavioral information obtained from bio-logging data using GPS-accelerometer tags provides understanding of how species shift strategies in response to environmental variability. This is vital to implement climate-adaptive conservation and management strategies. Given the growing availability of long-term accelerometer datasets within the ecological community, we recommend integrating such high-resolution information into conservation programs.

## Introduction

Planet Earth faces the dramatic and accelerating consequences of climate change with the expansion and intensification of anthropogenic stressors (Halpern *et al.* 2019; Abrahms *et al.* 2023). Ongoing global conservation and resource management efforts see the designation of protected areas as a widely used strategy to maintain species richness and population abundance and preserve ecosystems’ functioning (Shin *et al.* 2022). The design and implementation of protected areas have evolved to support resilient social-ecological systems (Padleton *et al.* 2017; Claudet *et al.* 2020). However, challenges affecting the effectiveness of protected areas include the continuous availability of data on current states and trends of nature and threats. Hence, long-term species monitoring programmes are essential to inform, implement and verify the effectiveness of such mitigation strategies (Soriano-Redondo *et al.* 2023).

Bio-logging technology, electronic devices attached to animals, provides vital insights into their movements and behavior, informing wildlife conservation and management efforts (Joly *et al.* 2019; Hindell *et al.* 2020; Davies *et al.* 2021). Population-level spatial distributions derived from bio-logging data has aided in assessing the extent and impact of human-wildlife interactions, such as interaction with fisheries and landscape fragmentation (Clay *et al.* 2019; Boudreau *et al.* 2022). However, animals use different areas for specific purposes like migration, foraging, or reproduction, emphasising each location’s unique functions in their life cycles. Hence it is important to identify how and why animals use specific areas, the relevance of these sites in face of long-term climatic shifts, and exposure to existing and emerging human pressures.

One of animals’ most important activities is finding and consuming food. Movement is necessary to forage for food and gain the energy needed to balance growth, body maintenance, reproduction, and physical activity (Pontzer & McGrosky 2022). Behavioural inference solely based on location data may misrepresent the underlying importance of the area used. Measures of daily physical activity, foraging intensity and efficiency can be used as indices to track spatio-temporal variations in energy spent and gained across the environment and/or resource exploitation. Accelerometer tags, record high-resolution data reflecting changes in speed due to the movement of the animal and provide measurements of effort and relative energy expenditure related to different behaviours and activities performed by individual animals, including foraging (Wilson *et al.* 2020; Chimienti *et al.* 2022). By recording at high frequency (> 1Hz), acceleration data capture behavioural activities not visible at coarser scale (e.g. prey hunting events) and, thus, are extremely useful for explaining area-use variation. When coupled with remote sensing data on environmental variability, accelerometer derived metrics can explain the animal decisions behind area-use variation and inform management and conservation.

The Bass Coast and Bass Strait waters are characterised by rich biodiversity, holding significant ecological, societal, and economic importance for both the region and the state of Victoria, Australia (Haddow *et al.* 2019). While a significant tourist and recreational destination, the region provides critical breeding habitat for charismatic marine top predators. Mainly, Phillip Island serves as a breeding area for approximately one-quarter of all seals in Victoria (McIntosh *et al.* 2018), approximately 1,000,000 short-tailed shearwaters (Latine name) visiting Australia, and the largest colony of Little Penguins (*Eudyptula minor*) consisting of ∼32,000 breeding adults (Sutherland & Dann 2012). Yet, the area is under climatic and anthropogenic pressure (Hobday & Pecl 2014; Halpern *et al.* 2019) threatening this globally Important Bird Area (IBA), and Marine Spatial Planning for the Bass Coast is currently under development. Overall, the Bass Strait’s ecological, economic, and historical significance underscores the need for its conservation and sustainable management to preserve its diverse marine life, support regional economies, and maintain its cultural heritage.

The long-term population monitoring programme at the penguins’ breeding colony of Phillip Island has regularly monitored individual penguins’ foraging behaviour during their breeding season with bio-logging technology, recording both horizontal and vertical movements. Little penguins are high trophic level generalist predators feeding in the shallower part of the water column (20-57 m), (Ropert-Coudert, Chiaradia & Kato 2006; Ropert-Coudert *et al.* 2006; Chiaradia *et al.* 2007) in dynamic coastal environments (Carroll *et al.* 2018; Meyer *et al.* 2020; Cavallo *et al.* 2020). Foraging performance (measured as trip duration, mass gain, and use of prey encounters) can vary within and among years, between adjacent colonies, as well as with environmental conditions, determining annual breeding success (Chiaradia & Nisbet 2006; Ropert-Coudert, Kato & Chiaradia 2009; Saraux *et al.* 2016; Sánchez *et al.* 2018; Joly *et al.* 2022).

Identifying and managing the areas where little penguins concentrate their foraging efforts is crucial to their survival and breeding performance. Although short-term studies have been conducted (Carroll *et al.* 2016; Sánchez *et al.* 2018), there is a lack of understanding of the long-term spatio-temporal variability of foraging intensity at sea, showing where, how and why foraging hotspots develop. Based on a ten-year bio-logging dataset on little penguins, we combine high-resolution bio-logging data from GPS-accelerometer loggers with metrics of environmental variability. Specifically, we test how the variation in Sea Surface Temperature (SST), thermocline, currents and water turbidity affect the foraging intensity of little penguins. These variables provide information on changes happening at lower trophic levels, affecting the abundance and distribution of prey consumed by little penguins. We predict different effects of these environmental parameters. Higher SST and stratified waters result in higher predictability of prey patches (Cox *et al.* 2018) leading to reduced foraging effort and increased efficiency (measured as average diving time and number of ondulations within dives over a foraging trip) (Meyer *et al.* 2020). However, increased SST can negatively affect penguins (Afán *et al.* 2015). Temperature above the 19-21 °C range can reduce foraging success (Carroll *et al.* 2016). Water turbidity might make prey more difficult to detect if these marine predators rely on visual cues for detecting and capturing their prey (Kowalczyk *et al.* 2015), as in shallow diving seabirds such as shearwaters (Darby *et al.* 2022). Overall, how these environmental conditions drive the spatio-temporal variability in foraging intensity across breeding seasons and stages is not known. Finally, given the current discussion to implement Marine Spatial Planning for the State of Victoria, Australia, we highlight the need for high resolution data to responsible stewardship and conservation off Phillip Island to protect this valuable marine ecosystem into the future.

## Material and Methods

### Penguin tracking and data preparation

This study covers 11 breeding seasons, from 2010 to 2020 (inclusive). During breeding (Austral spring/summer, from August to February), the little penguin is a central-place forager with foraging trips between 1 and 16 days (Saraux *et al.* 2011). Typically, their breeding season can be divided into three stages: *Incubation* (lasting ∼35 days), *Guard* (∼2 weeks) during which both parents alternate foraging trips at sea and *Post Guard* (∼5-8 weeks), when older chicks are left alone during the day and parents come back at night for feeding (Chiaradia & Nisbet 2006). Penguins were captured from their nest boxes and equipped with GPS-Accelerometer loggers attached to the lower dorsal region of the bird using Tesa® tape.

Given the length of the study and the rapid developments in bio-logging technology within these ten years, data were collected using different loggers (Little Leonardo, Japan, TechnoSmart, Italy and WACU IPHC Strasbourg for accelerometer loggers; and Cat Track, and Technosmart for GPS loggers, and at varying sampling resolutions depending on logger manufacture, battery and storage capacity, animals’ breeding stage and length of the foraging trips (Zimmer *et al.* 2011; Sánchez *et al.* 2018; Dupuis *et al.* 2023). Loggers were set to record: i) GPS locations at an interval from 10 sec to 5 min, depending on logger type, season and breeding stage (Dupuis *et al.* 2023), ii) underwater pressure (in millibars) or depth (in meter) at 1 Hz and iii) tri-axial acceleration at 25 or 50 Hz. As penguins do not forage at night, only GPS data between nautical dawn and dusk were analysed, generating diurnal “*foraging segments”*. GPS locations for each foraging segment were checked, filtered, and interpolated at 15 min intervals (Dupuis *et al.* 2023). Step length was then calculated from interpolated foraging segments using the R package *MomentuHMM* (McClintock & Michelot 2018). Underwater pressure was converted in meters of depth and tri-axial acceleration data were subsampled at 25 Hz. Both depth and acceleration data were treated and analysed as in Chimienti *et al.* 2022. Specifically, the approach using the supervised machine learning algorithm, *Random Forest,* developed in Chimienti *et al.* 2022 was used to infer the behaviours little penguins performed while foraging at sea, such as hunting, swimming, descending and ascending within the water column, porpoising and preening near the water surface (Chimienti *et al.* 2022). GPS locations, diving depth, and resulting information from behavioural classification were then matched by date and time. Since many dives could be performed within the 15 minutes between the two consecutive GPS locations, for each time interval we calculated the average maximum diving depth and time spent hunting indicating hunting effort.

### Environmental data

Data on bathymetry, thermocline and currents were used as in Sanchez *et al.* 2019 and Joly *et al.* 2022. Specifically, we used depth (m) and intensity (°C) of thermocline and currents’ speed (m/s). Sea Surface Temperature (SST, °C) and Secchi Disk Depth (ZSD, m) were downloaded from the Copernicus database gap-free L4 products (https://www.copernicus.eu/en/copernicus-services/marine). ZSD was used as a metric of light transmissibility through the water column, i.e. turbidity. Greater ZSD corresponds to clearer water. All variables were recorded daily at variable resolutions (from 250x250 m for bathymetry up to 10x10 Km for the thermocline and currents). Hence all variables were rasterised at 10x10 Km grid.

### Data analysis

The analysis aims at quantifying the drivers of the spatio-temporal variability in the foraging intensity of little penguins across breeding seasons and stages. Hence only GPS locations where penguins were diving were considered for further analyses. Given the spatio-temporal discrepancy between the small movement scale of the little penguins (Figure S1, *Supporting Information*) and the scale of the available environmental data, the combined GPS dataset was rasterised at the same resolution as the environmental data. Daily rasters were calculated at 10x10 km considering the reproductive stage and recording for each cell the median values for time spent hunting, maximum dive depth and step length, number of data points and number of unique individuals. These datasets were then matched with the environmental variables. SST values were averaged over weekly periods, given that mismatches between SST and the distribution of predators suggest time lags effects (Grémillet *et al.* 2008; Ramírez *et al.* 2016).

We used Generalized Additive Mixed Models (GAMMs, *mgcv* package, (Wood 2011) to investigate the relationships between foraging intensity, animals’ movement characteristics and environmental conditions. Time spent hunting was used as a response variable, while maximum diving depth, step length, bathymetry, thermocline depth and gradient, currents’ speed, SST and ZSD were included as explanatory variables. Spatial coordinates were added to the model to account for spatial autocorrelation, breeding stage and season were added as fixed effects since we were interested in the between years comparison. Before modelling, all explanatory variables were checked for multi-collinearity. Thermocline depth and gradient were modelled within the same spline to quantify the effect of the thermocline conditions on foraging intensity. We performed model selection on the full model for the type of spline to use; isotropic (*s*) versus the anisotropic (*te, ti*) splines. Data exploration showed variation in SST values across breeding stages (see *Results*). Hence, we hypothesised that the relationships with SST values were likely to change between breeding stages, and we used the “*by*” term within the spline for SST adding the interaction with the breeding stage. *Day* was used as grouping for random effects. Starting from the full model structure, we run models with all possible variable combinations using the “MuMin” package . The model with the lowest corrected Akaike Information Criterion (AICc) was selected as best, and we reported and considered the top 5 models as competing models. Models were run using the *Tweedie* distribution family and checked for violations of model assumptions in terms of residual autocorrelation, heterogeneity, and normality. All data manipulation and analysis were performed in R statistical computing software version 4.2.2, (R Core Team 2022).

## Results

We obtained 247 individual tracks across 11 breeding seasons. Penguins were sampled once within the same breeding season. In contrast, 36% of the individuals were sampled more than once across all seasons (2-5 times, with the majority being 2 or 3). GPS-Accelerometer data were available for *Guard* stage for the 2010-2020 period. *Incubation* and *Post Guard* were obtained from 2014 and 2015, respectively (Table 1). The number of individual penguins tracked and data points varied across seasons (Table 1) as well as the timeframe when tracking occurred during the season, temporal overlap (Figure S2) and spatial distribution (Figure S3, S4).

**Table 1:**
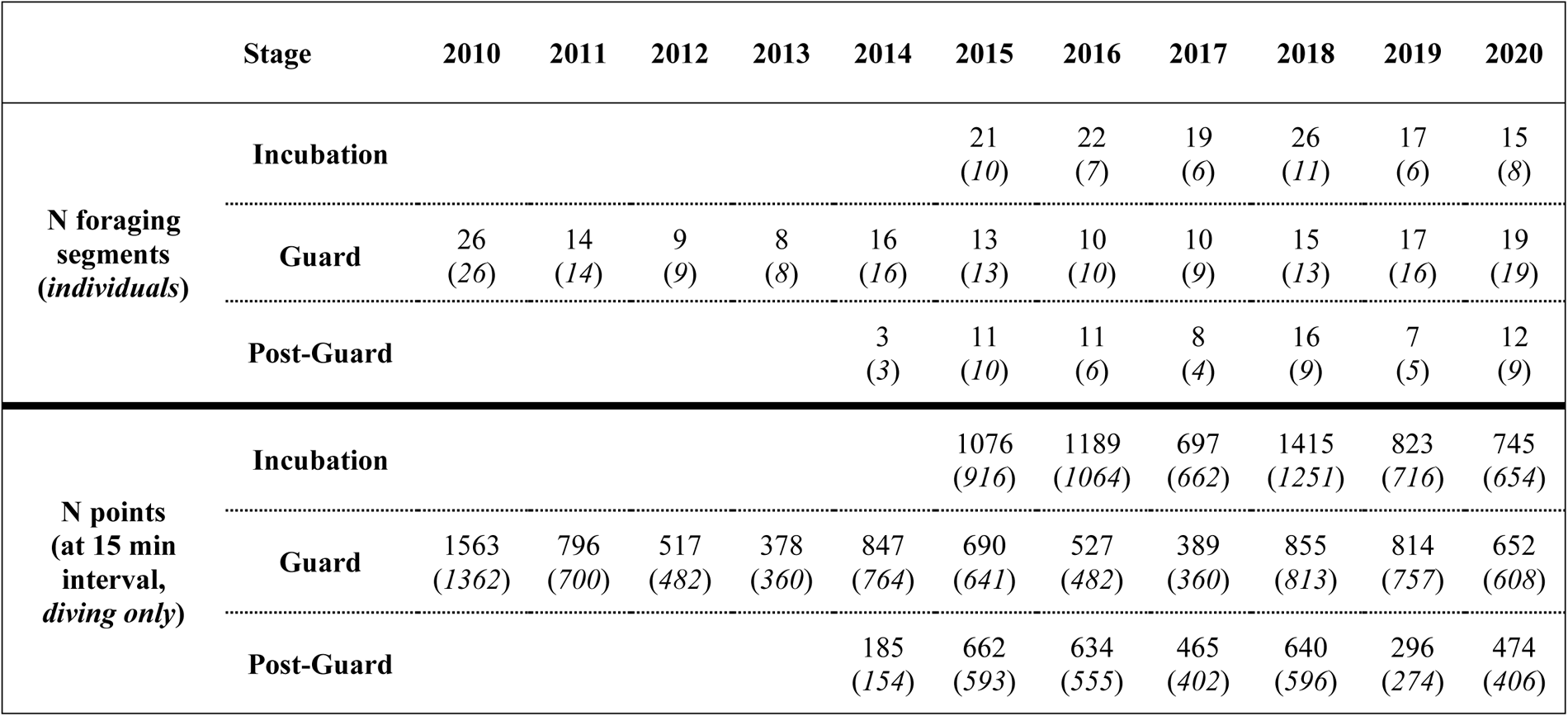
Overview of data collected using GPS-Accelerometer bio-logging tags on Little penguin (*Eudyptula minor*) during the breeding seasons 2010-2020 and across the three stages: Incubation, guard, post guard. 36% of these individuals have been tracked from 2-5 times.

Time spent hunting (sec) associated with the 15 min GPS locations showed variation between seasons (Figure 1 and S5, mean ± sd from 2010 to 2020 respectively: 38.4 ± 36.5, 45.0 ± 45.4, 42.96 ± 33.6, 111.91 ± 83.93, 41.2 ± 35.9, 60.8 ± 47.9, 54.6 ± 53.3, 44.6 ± 38.8, 81.9 ± 67.7, 39.3 ± 32.7) and consistency between stages (time spent hunting, mean ± sd: *Incubation* 51.8 ± 50.3, *Guard* 50.8 ± 47.7, *Post Guard* 53.8 ± 50.2). Across all seasons, individuals hunted within the upper part of the water column (Figure S5, mean ± sd for all seasons, 22.7 ± 10.8 m), with 2013 being the only season when penguins performed shallower dives compared to the other years (Figure S5). Step lengths recorded across seasons and stages were also highly overlapping (Figure S6 and S7, mean ± sd for all seasons 791.5 ± 358.2 m).

**Figure 1:**
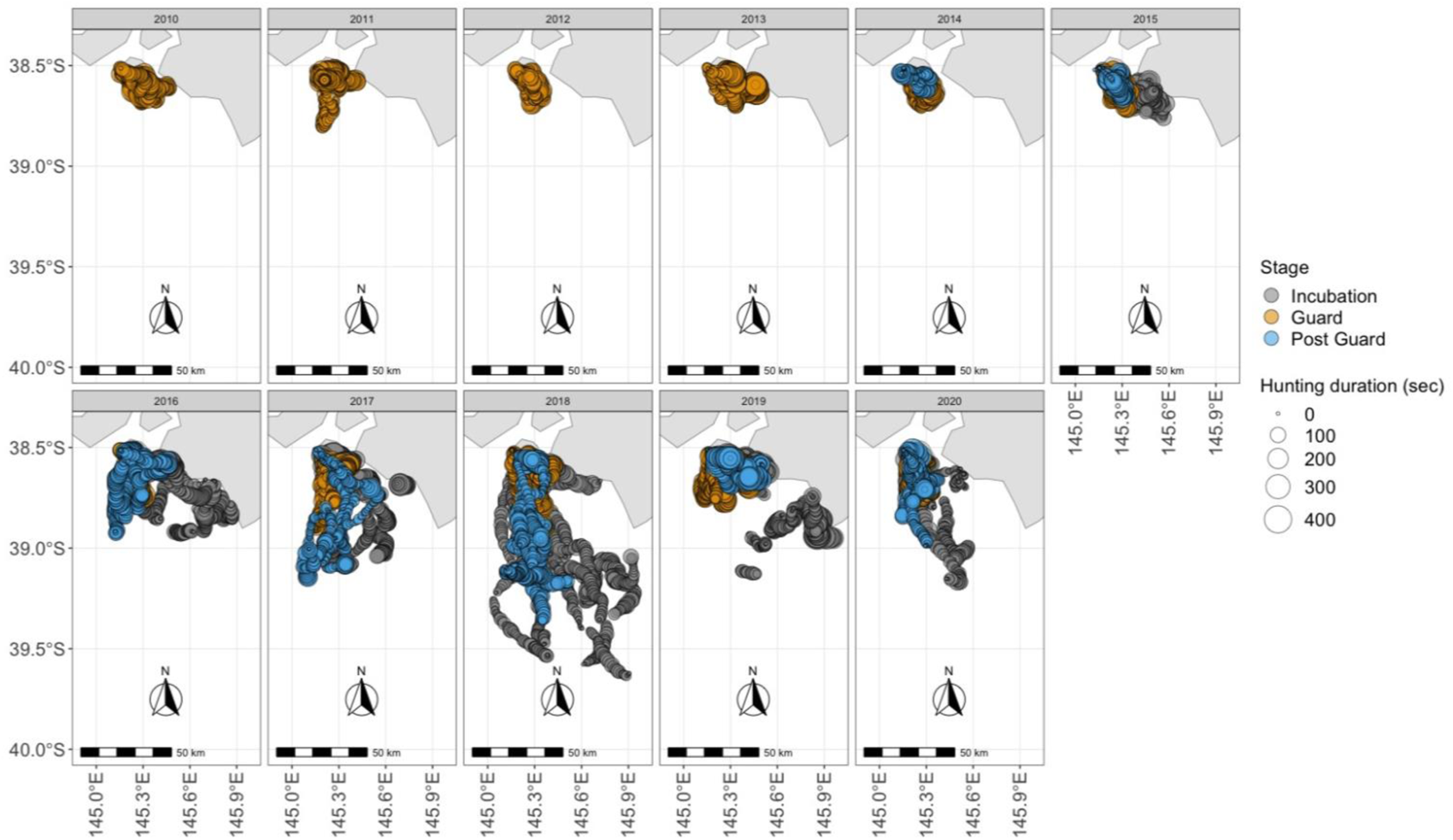
Map of the study area showing the overview of the GPS-Accelerometer dataset collected on Little penguin (*Eudyptula minor*) across eleven sampling seasons. Colors indicate breeding stages: *Incubation* (grey), *Guard* (gold) and *Post Guard* (blue). Size indicates time spent hunting (sec).

Hunting areas had shallow and weak thermoclines and slow currents with little to no variation across years and breeding stages (Figure S6). The bathymetric profiles of the area also overlapped across seasons and stages (mean ± sd for all seasons 57.04 ± 18.2 m). Over the seasons, little penguins used areas with variable turbidity levels (Figure S6 and S7, from 2010 to 2020 respectively 13.2 ± 3.1, 17.8 ± 4.08, 16.7 ± 2.5, 11.9 ± 3.9, 14.05 ± 2.9, 18.4 ± 3.7, 10.5 ± 3.2, 19.7 ± 4.7, 23.2 ± 4.6, 16.4 ± 4.0, 15.6 ± 4.13 m). SST varied between seasons and increased with the progress of the breeding seasons (mean ± sd 13.8 ± 0.8, 14.9 ± 1.37, 15.4 ± 1.6 °C), specifically during *Guard* and *Post Guard* stages (Figure S6 and S7).

The full GAM model with tensor product spline *te* combining the thermocline depth and gradient had the lowest AIC values compared to the other splines (10464.62, 10465.20, 10462.07 and 10463.81, respectively, for models with the thermocline variables modelled in two separate splines, or together within the isotropic *s* spline, the anisotropic *te* and *ti* splines). Model selection for the model testing the variation in hunting time across years and stages as a function of environmental variability, favoured the formula including maximum diving depth, SST, step length, ZSD, stage and season as fixed effects (Table S1, R-sq. adj = 0.485). The model detected seasonal variation in hunting intensity, with higher levels during the *Post Guard* stage (p-value <0.05, Table S2). Time spent hunting followed a sigmoidal behaviour in relation to maximum diving depth (p-value <0.001, Figure 2). Time spent hunting steeply increased up to 10 m in depth, after which it increased very slowly for deeper depths. Increased hunting time was associated with shorter step lengths (p-value <0.001, Figure 2) and with higher values of water turbidity (ZSD, p-values <0.01, Figure 2). Higher SST values had a negative effect on hunting time during *Guard* and *Post Guard* stages (p-values <0.001, Figure 3) but not in *Incubation*. At the spatial level, penguins hunted mostly on the coastal areas on the east side of the breeding colony off the Kilcunda Reserve, with some individuals reaching the farthest south-west areas (Figure 4).

**Figure 2:**
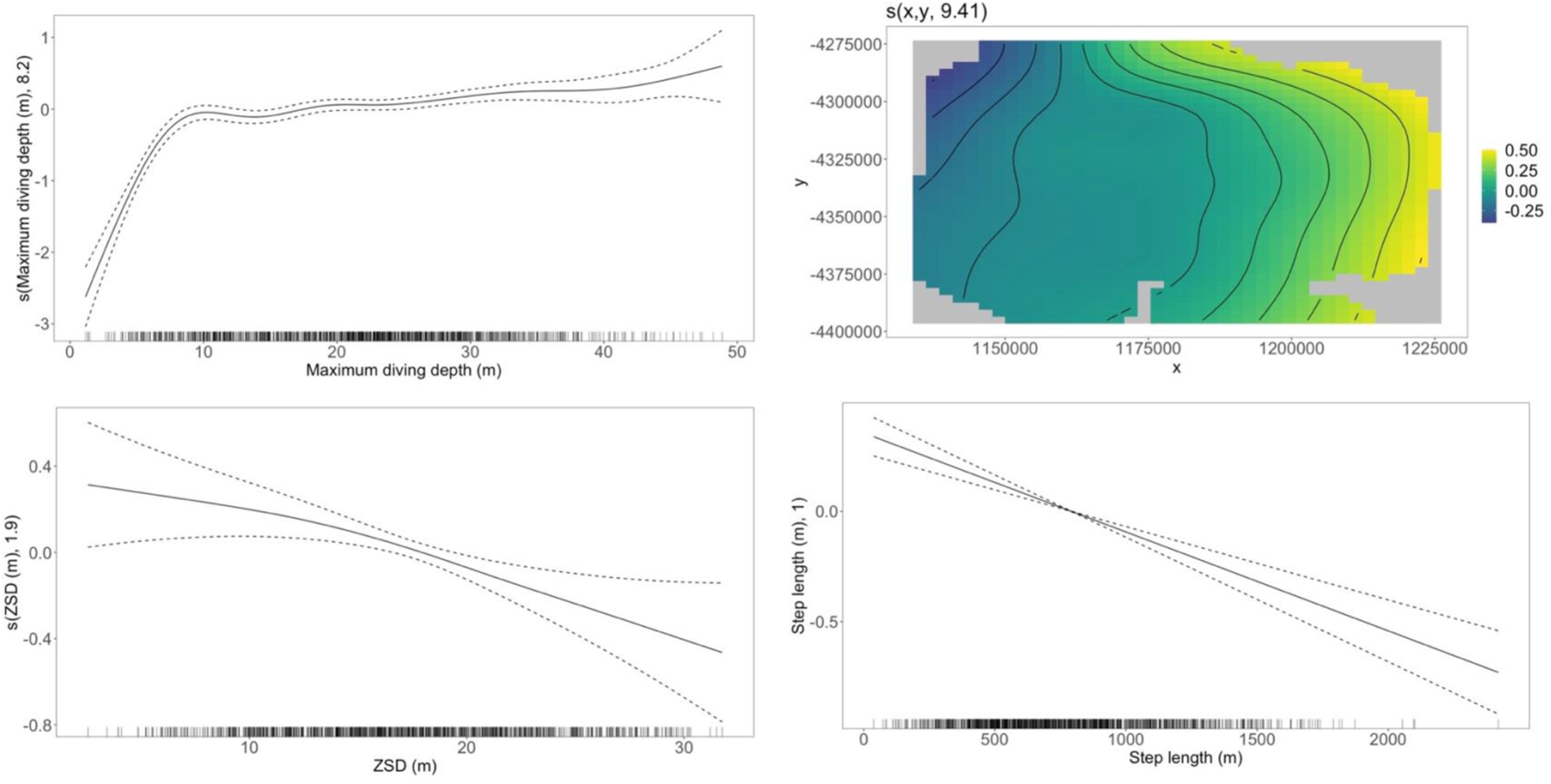
Significant GAM covariates describing the time spent hunting in Little penguins (*Eudyptula minor*) as a function of maximum diving depth (m), spatial usage, Secchi disk depth (ZSD)/water turbidity (m), and step length (m).

**Figure 3:**
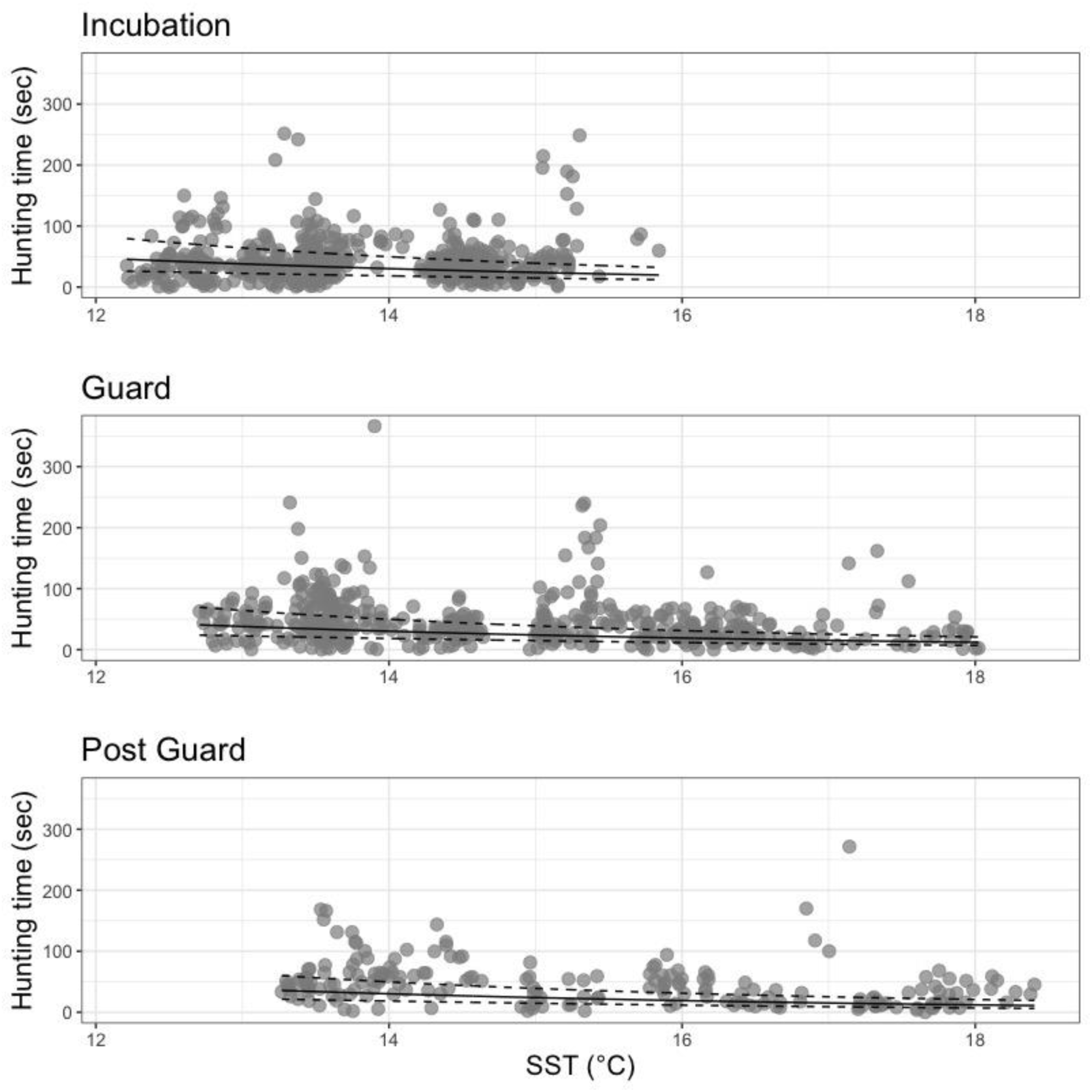
Time spent hunting (sec) as function of Sea Surface Temperature (SST, °C) by breeding stage, *Incubation*, *Guard* and *Post Guard*. Solid and dashed lines indicate the predictions and confidence intervals from the Generalized Additive Model.

**Figure 4:**
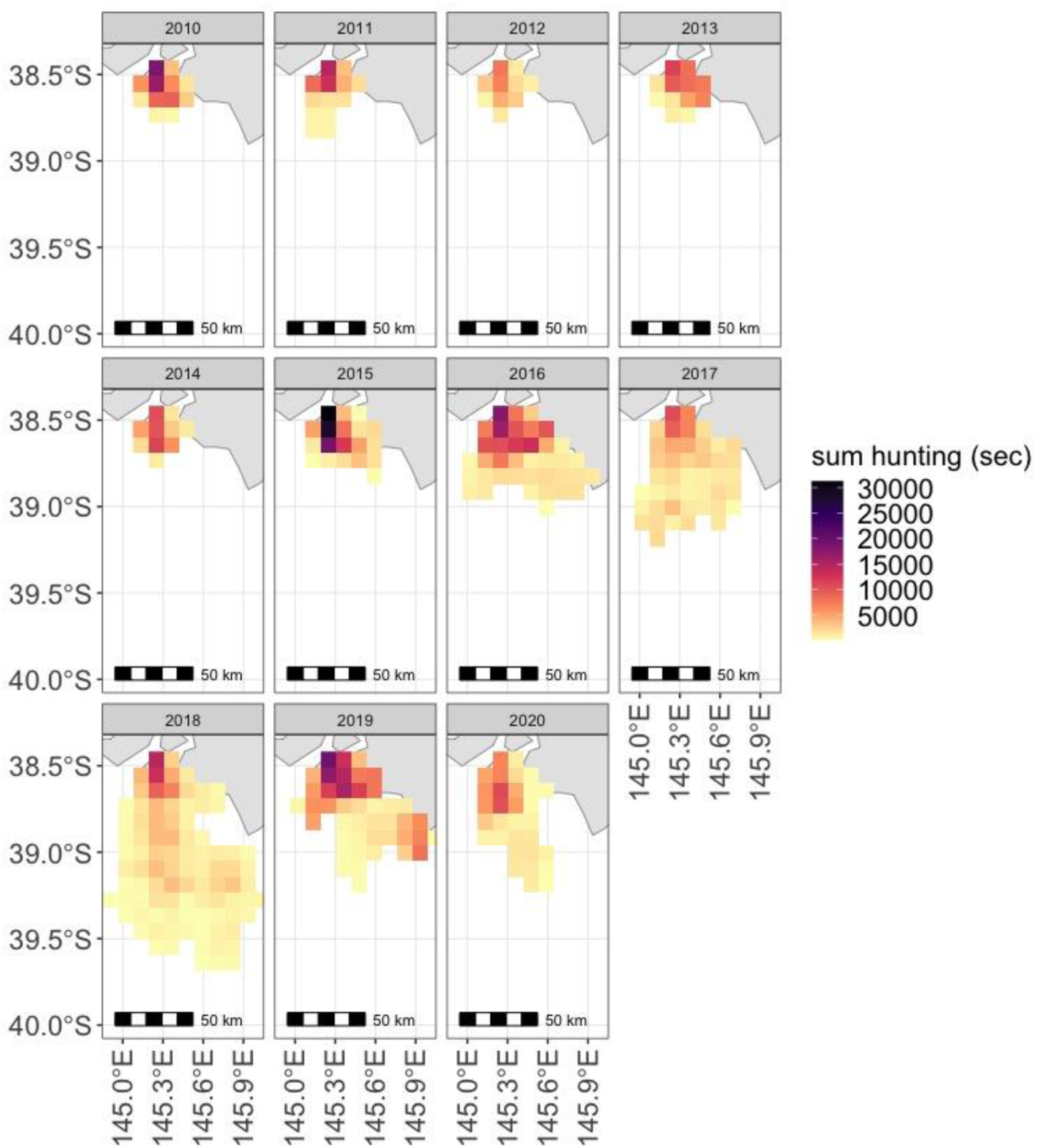
Total time spent hunting, obtained from GPS-accelerometer datasets, performed by Little penguin (*Eudyptula minor*) across eleven breeding seasons. Rasters are at 10x10 Km resolution.

## Discussion

We used a unique long-term (ten years) high-resolution bio-logging dataset to quantify how, where, and why little penguins breeding on Phillip Island invest in their hunting activity in relation to environmental conditions. We used satellite derived environmental paramters indicating horizontal variability of surface waters (e.g. SST) as well as the vertical stratification of the water (e.g. thermocline depth, water turbidity). Hence, we assessed the species’ capacity to adjust their behaviour in both horizontal and vertical space. Penguins increased their time spent hunting in coastal, colder and more turbid waters while reducing spatial displacement (shorter step lengths) and diving deeper; with a slower increase in time spent hunting below 10 m of depth (max diving depth 74 m, a new record for the species – see Ropert-Coudert *et al.* 2006). Horizontal and vertical movements reveal distinct facets of behavior; horizontal movements may respond to large-scale general environmental cues such as SST, while vertical movements may respond to more localised prey presences and prey pursuit tactics (Wilson, Ropert-Coudert & Kato 2002; Ropert-Coudert *et al.* 2006; Ramasco *et al.* 2015). While sample sizes were unbalanced among years and relatively small in some years, due to the combination of GPS and accelerometer data, the great inter-annual variation in little penguin’s spatial distribution was further confirmed by the analysis of core and home ranges (50% and 95% UD) over the same period based on the more robust and consistent GPS-only dataset (Dupuis *et al.* 2023).

Productive aquatic ecosystems can be characterised by turbid waters (Abrahams & Kattenfeld 1997). High turbidity in coastal waters can also result from human-induced activities, freshwater inflow, or wind-driven resuspension of sediments (Arendt *et al.* 2011; Ortega *et al.* 2020), which is known to enhance productivity (Kowalczyk *et al.* 2015). Marine predators, as little penguins, can use various sensory cues to detect and hunt their prey in turbid or dark waters (Davis *et al.* 1999; Darby *et al.* 2022). The effect of water visibility/turbidity on prey detection and foraging in aquatic systems supports contrasting theories depending on foraging strategies and type of prey hunted. High levels of turbidity can influence predator-prey interactions by lowering prey capture and offering refuge from predation but also elevate predation by enhancing contrast and providing abundant resources (Abrahams & Kattenfeld 1997; Ortega *et al.* 2020; Goździejewska & Kruk 2022). Hence, it is possible that, in turbid waters, penguins either exploit the increased availability of nutrients acquiring more prey or require longer time to locate and catch their prey. The effects of climate and human driven habitat changes on water turbidity and how they affect trophic interactions remain understudied (Goździejewska & Kruk 2022). Further work should be directed towards how this species benefits from highly turbid areas and the related foraging success.

An increase in SST, marine heatwaves and changes in the predominance of the main currents influencing the dynamics of the Bass Strait (the nutrient-poor South Australian Current, the nutrient-rich sub-Antarctic Surface Water and the nutrient-poor East Australian Current) are likely also to shift the trophic interactions within the area (Poloczanska *et al.* 2007; Afán *et al.* 2015; Oliver *et al.* 2017). SST distributions within the little penguins’ foraging area varied between years. Additionally, little penguins are exposed to warmer waters as the breeding season progresses from spring to summer. Our analysis showed that hunting time decreased with warmer waters, especially during the *Guard* and *Post Guard* stages (min-max SST 12-18.4 °C). During the breeding season, little penguins can switch foraging strategy, requiring different hunting time, and targeting different types of prey and prey aggregations with varying energetic content (Cavallo *et al.* 2020; Sutton & Arnould 2022). More dispersed and/or less abundant prey in colder and less-stratified waters, or increased prey availability in warmer and stratified waters (Meyer *et al.* 2020) can lead to changes in effort allocated to hunting activities. An increase in SST beyond the range measured in this study is happening faster and more strongly than anticipated, reducing foraging success (Carroll *et al.* 2016) and negatively affecting breeding success and mean laying date (Pulvirenti, Reina & Chiaradia 2023).

Environmental variables’ relatively coarse spatial and temporal resolution may account for some unexplained variation in our models, along with other unknown factors, such as vertical prey distribution and true profitability, ie, prey capture success. Our estimate of time spent hunting does not necessarily indicate a greater prey intake (Florko *et al.* 2023). During breeding, little penguins must return to the colony to either attend the nest or feed for the chicks. Hence, prey might differ when feeding for chicks vs. for themselves, due to different energy requirements, higher need of proteins for chicks (Chiaradia *et al.* 2016). While little penguins maximise foraging performance by concentrating efforts on larger quantities of prey (Sutton & Arnould 2022), high nutritious contents (Chiaradia *et al.* 2012) and lower trophic levels (Chiaradia *et al.* 2010), further investigation requires validating links between high-resolution prey availability, prey energy content, energy expenditure and hunting intensity. Changing environments, shifts across trophic levels as well as in energy balances (energy gain versus energy expenditure) can ultimately affect breeding performance (Joly *et al.* 2022) and life history strategies (Pontzer & McGrosky 2022).

### Policy Implications

South-east Australia is experiencing significant changes in oceanographic features and species distributions (Wu et al. 2012; Hobday & Pecl 2014), with the ocean temperature increasing twice as fast as the global average (CSIRO and Bureau of Meteorology 2022). The threatening processes affecting marine biodiversity are related to climate change and human activities. Ocean current shifts, sea level changes, pH fluctuations, temperature variations, and increased storms will likely disrupt various ecological processes, leading to reduced species availability, habitat areas, and breeding and foraging sites. While Tourism doesn’t affect Phillip Island’s little penguins, human activities at sea, such as maritime traffic, have negatively affected foraging efficiency (measured as mass gained at sea, Dupuis *et al.* 2023). Our analysis reveals little penguins’ habitat use variation in the Bass Strait across environmental conditions and breeding stages, emphasizing the vital role of SST and highly productive coastal areas in sustaining their foraging activities.

Integrating climate change into Marine Protected Area (MPA) management is crucial (Ramírez *et al*. 2017; Zentner *et al*. 2023). By analyzing long-term high-resolution GPS-accelerometer data, we discovered the adaptable foraging strategies of these marine predators. This knowledge equips managers with precise insights, enabling effective conservation planning in response to environmental shifts. Understanding animal responses to fluctuating conditions is vital for predicting future behavior. Our decade-long analysis provides extensive spatial and temporal replication, empowering informed decision-making for the Bass Strait’s conservation initiatives.

## Author contributions

Conceptualisation: MC, YRC, AK, AC, Analysis: MC, BD and NJ respectively provided GPS data and environmental data analysed. Writing: all authors with MC leading, Review of draft: all authors. Supervision : YRC, AK, AC

## Data availability statement

R codes used to manipulate and analyse the data used are available at https://github.com/MariannaChimi/MuFFIN_MSCA_II. Data will be made available after acceptance.

## Acknowledgements

We thank Phillip Island Nature Parks for their continued support and commitment to penguin research. We thank all the students and volunteers for their tireless support in collecting these data over the years, particularly the Nature Parks’s research technical staff Leanne Renwick, Paula Wasiak, Meagan Tucker, Marjolein van Polanen Petel and Jordan Roberts. Field work protocol was approved by the ethics committee of the Phillip Island Nature Park Animal Experimentation Committee with a research permit issued by the Department of Environment, Land, Water and Planning of Victoria, Australia.

## References

1. Abrahams, M. & Kattenfeld, M. (1997) The role of turbidity as a constraint on predator-prey interactions in aquatic environments. Behavioral Ecology and Sociobiology, 40, 169– 174.

2. Abrahms, B., Carter, N.H., Clark-Wolf, T.J., Gaynor, K.M., Johansson, E., McInturff, A., Nisi, A.C., Rafiq, K. & West, L. (2023) Climate change as a global amplifier of human–wildlife conflict. Nature Climate Change.

3. Afán, I., Chiaradia, A., Forero, M.G., Dann, P. & Ramírez, F. (2015) A novel spatio-temporal scale based on ocean currents unravels environmental drivers of reproductive timing in a marine predator. Proceedings. Biological Sciences / the Royal Society, 282.

4. Arendt, K.E., Dutz, J., Jonasdottir, S.H., Jung-Madsen, S., Mortensen, J., Moller, E.F. & Nielsen, T.G. (2011) Effects of suspended sediments on copepods feeding in a glacial influenced sub-Arctic fjord. Journal of Plankton Research, 33, 1526–1537.

5. Boudreau, M.R., Gantchoff, M.G., Ramirez-Reyes, C., Conlee, L., Belant, J.L. & Iglay, R.B. (2022) Using habitat suitability and landscape connectivity in the spatial prioritization of public outreach and management during carnivore recolonization. Journal of Applied Ecology, 59, 757–767.

6. Carroll, G., Everett, J.D., Harcourt, R., Slip, D. & Jonsen, I. (2016) High sea surface temperatures driven by a strengthening current reduce foraging success by penguins. Scientific Reports, 6, 22236.

7. Carroll, G., Harcourt, R., Pitcher, B.J., Slip, D. & Jonsen, I. (2018) Recent prey capture experience and dynamic habitat quality mediate short-term foraging site fidelity in a seabird. Proceedings. Biological Sciences / the Royal Society, 285.

8. Cavallo, C., Chiaradia, A., Deagle, B.E., Hays, G.C., Jarman, S., McInnes, J.C., Ropert-Coudert, Y., Sánchez, S. & Reina, R.D. (2020) Quantifying prey availability using the foraging plasticity of a marine predator, the little penguin. Functional Ecology.

9. Chiaradia, A., Forero, M.G., Hobson, K.A. & Cullen, J.M. (2010) Changes in diet and trophic position of a top predator 10 years after a mass mortality of a key prey. ICES Journal of Marine Science, 67, 1710–1720.

10. Chiaradia, A., Forero, M.G., Hobson, K.A., Swearer, S.E., Hume, F., Renwick, L. & Dann, P. (2012) Diet segregation between two colonies of little penguins Eudyptula minor in southeast Australia. Austral ecology, 37, 610–619.

11. Chiaradia, A. & Nisbet, I.C.T. (2006) Plasticity in parental provisioning and chick growth in Little Penguins *Eudyptula minor* in years of high and low breeding success . Ardea, 94, 257–270.

12. Chiaradia, A., Ramírez, F., Forero, M.G. & Hobson, K.A. (2016) Stable Isotopes (δ13C, δ15N) Combined with Conventional Dietary Approaches Reveal Plasticity in Central-Place Foraging Behavior of Little Penguins Eudyptula minor. Frontiers in ecology and evolution, 3.

13. Chiaradia, A., Ropert-Coudert, Y., Kato, A., Mattern, T. & Yorke, J. (2007) Diving behaviour of Little Penguins from four colonies across their whole distribution range: bathymetry affecting diving effort and fledging success. Marine Biology, 151, 1535– 1542.

14. Chimienti, M., Kato, A., Hicks, O., Angelier, F., Beaulieu, M., Ouled-Cheikh, J., Marciau, C., Raclot, T., Tucker, M., Wisniewska, D.M., Chiaradia, A. & Ropert-Coudert, Y. (2022) The role of individual variability on the predictive performance of machine learning applied to large bio-logging datasets. Scientific Reports, 12, 19737.

15. Claudet, J., Loiseau, C., Sostres, M. & Zupan, M. (2020) Underprotected marine protected areas in a global biodiversity hotspot. One Earth, 2, 380–384.

16. Clay, T.A., Small, C., Tuck, G.N., Pardo, D., Carneiro, A.P.B., Wood, A.G., Croxall, J.P., Crossin, G.T. & Phillips, R.A. (2019) A comprehensive large-scale assessment of fisheries bycatch risk to threatened seabird populations. Journal of Applied Ecology.

17. Cox, S.L., Embling, C.B., Hosegood, P.J., Votier, S.C. & Ingram, S.N. (2018) Oceanographic drivers of marine mammal and seabird habitat-use across shelf-seas: A guide to key features and recommendations for future research and conservation management. Estuarine, Coastal and Shelf Science, 212, 294–310.

18. CSIRO and Bureau of Meteorology. (2022) State of the Climate 2022. Commonwealth of Australia 2023, Bureau of Meteorology (ABN 92 637 533 532), CRICOS Provider 02015K.

19. Darby, J., Clairbaux, M., Bennison, A., Quinn, J.L. & Jessopp, M.J. (2022) Underwater visibility constrains the foraging behaviour of a diving pelagic seabird. Proceedings. Biological Sciences / the Royal Society, 289, 20220862.

20. Davies, T.E., Carneiro, A.P.B., Tarzia, M., Wakefield, E., Hennicke, J.C., Frederiksen, M., Hansen, E.S., Campos, B., Hazin, C., Lascelles, B., Anker-Nilssen, T., Arnardóttir, H., Barrett, R.T., Biscoito, M., Bollache, L., Boulinier, T., Catry, P., Ceia, F.R., Chastel, O., Christensen-Dalsgaard, S., Cruz-Flores, M., Danielsen, J., Daunt, F., Dunn, E., Egevang, C., Fagundes, A.I., Fayet, A.L., Fort, J., Furness, R.W., Gilg, O., González-Solís, J., Granadeiro, J.P., Grémillet, D., Guilford, T., Hanssen, S.A., Harris, M.P., Hedd, A., Huffeldt, N.P., Jessopp, M., Kolbeinsson, Y., Krietsch, J., Lang, J., Linnebjerg, J.F., Lorentsen, S., Madeiros, J., Magnusdottir, E., Mallory, M.L., McFarlane Tranquilla, L., Merkel, F.R., Militão, T., Moe, B., Montevecchi, W.A., Morera-Pujol, V., Mosbech, A., Neves, V., Newell, M.A., Olsen, B., Paiva, V.H., Peter, H., Petersen, A., Phillips, R.A., Ramírez, I., Ramos, J.A., Ramos, R., Ronconi, R.A., Ryan, P.G., Schmidt, N.M., Sigurðsson, I.A., Sittler, B., Steen, H., Stenhouse, I.J., Strøm, H., Systad, G.H.R., Thompson, P., Thórarinsson, T.L., Bemmelen, R.S.A., Wanless, S., Zino, F. & Dias, M.P. (2021) Multispecies tracking reveals a major seabird hotspot in the North Atlantic. Conservation letters.

21. Davis, R.W., Fuiman, L.A., Williams, T.M., Collier, S.O., Hagey, W.P., Kanatous, S.B., Kohin, S. & Horning, M. (1999) Hunting behavior of a marine mammal beneath the antarctic fast Ice. Science, 283, 993–996.

22. Dupuis, B., Kato, A., Joly, N., Saraux, C., Ropert-Coudert, Y., Chiaradia, A. & Chimienti, M. (2023) COVID-related anthropause highlights the impact of marine traffic on breeding little penguins. BioRxiv.

23. Florko, K.R.N., Shuert, C.R., Cheung, W.W.L., Ferguson, S.H., Jonsen, I.D., Rosen, D.A.S., Sumaila, U.R., Tai, T.C., Yurkowski, D.J. & Auger-Méthé, M. (2023) Linking movement and dive data to prey distribution models: new insights in foraging behaviour and potential pitfalls of movement analyses. Movement Ecology, 11, 17.

24. Goździejewska, A.M. & Kruk, M. (2022) Zooplankton network conditioned by turbidity gradient in small anthropogenic reservoirs. Scientific Reports, 12, 3938.

25. Grémillet, D., Lewis, S., Drapeau, L., van Der Lingen, C.D., Huggett, J.A., Coetzee, J.C., Verheye, H.M., Daunt, F., Wanless, S. & Ryan, P.G. (2008) Spatial match-mismatch in the Benguela upwelling zone: should we expect chlorophyll and sea-surface temperature to predict marine predator distributions? Journal of Applied Ecology, 45, 610–621.

26. Haddow, J., Duncan, J., Kilborn, A., Meredith, C. & Wescott, G. (2019) Assessment of the Values of Victoria’s Marine Environment . Victorian Environmental Assessment Council.

27. Halpern, B.S., Frazier, M., Afflerbach, J., Lowndes, J.S., Micheli, F., O’Hara, C., Scarborough, C. & Selkoe, K.A. (2019) Recent pace of change in human impact on the world’s ocean. Scientific Reports, 9, 11609.

28. Hindell, M.A., Reisinger, R.R., Ropert-Coudert, Y., Hückstädt, L.A., Trathan, P.N., Bornemann, H., Charrassin, J.-B., Chown, S.L., Costa, D.P., Danis, B., Lea, M.-A., Thompson, D., Torres, L.G., Van de Putte, A.P., Alderman, R., Andrews-Goff, V., Arthur, B., Ballard, G., Bengtson, J., Bester, M.N., Blix, A.S., Boehme, L., Bost, C.-A., Boveng, P., Cleeland, J., Constantine, R., Corney, S., Crawford, R.J.M., Dalla Rosa, L., de Bruyn, P.J.N., Delord, K., Descamps, S., Double, M., Emmerson, L., Fedak, M., Friedlaender, A., Gales, N., Goebel, M.E., Goetz, K.T., Guinet, C., Goldsworthy, S.D., Harcourt, R., Hinke, J.T., Jerosch, K., Kato, A., Kerry, K.R., Kirkwood, R., Kooyman, G.L., Kovacs, K.M., Lawton, K., Lowther, A.D., Lydersen, C., Lyver, P.O., Makhado, A.B., Márquez, M.E.I., McDonald, B.I., McMahon, C.R., Muelbert, M., Nachtsheim, D., Nicholls, K.W., Nordøy, E.S., Olmastroni, S., Phillips, R.A., Pistorius, P., Plötz, J., Pütz, K., Ratcliffe, N., Ryan, P.G., Santos, M., Southwell, C., Staniland, I., Takahashi, A., Tarroux, A., Trivelpiece, W., Wakefield, E., Weimerskirch, H., Wienecke, B., Xavier, J.C., Wotherspoon, S., Jonsen, I.D. & Raymond, B. (2020) Tracking of marine predators to protect Southern Ocean ecosystems. Nature, 580, 87–92.

29. Hobday, A.J. & Pecl, G.T. (2014) Identification of global marine hotspots: sentinels for change and vanguards for adaptation action. Reviews in Fish Biology and Fisheries, 24, 415–425.

30. Joly, N.B., Chiaradia, A., Georges, J.Y. & Saraux, C. (2022) Environmental effects on foraging performance in little penguins: a matter of phenology and short-term variability. Marine Ecology Progress Series, 692, 151–168.

31. Joly, K., Gurarie, E., Sorum, M.S., Kaczensky, P., Cameron, M.D., Jakes, A.F., Borg, B.L., Nandintsetseg, D., Hopcraft, J.G.C., Buuveibaatar, B., Jones, P.F., Mueller, T., Walzer, C., Olson, K.A., Payne, J.C., Yadamsuren, A. & Hebblewhite, M. (2019) Longest terrestrial migrations and movements around the world. Scientific Reports, 9, 15333.

32. Kowalczyk, N.D., Reina, R.D., Preston, T.J. & Chiaradia, A. (2015) Selective foraging within estuarine plume fronts by an inshore resident seabird. Frontiers in Marine Science, 2.

33. McClintock, B.T. & Michelot, T. (2018) momentuHMM :R package for generalized hidden Markov models of animal movement. Methods in Ecology and Evolution, 9, 1518– 1530.

34. McIntosh, R.R., Kirkman, S.P., Thalmann, S., Sutherland, D.R., Mitchell, A., Arnould, J.P.Y., Salton, M., Slip, D.J., Dann, P. & Kirkwood, R. (2018) Understanding meta-population trends of the Australian fur seal, with insights for adaptive monitoring. Plos One, 13, e0200253.

35. Meyer, X., MacIntosh, A.J.J., Chiaradia, A., Kato, A., Ramírez, F., Sueur, C. & Ropert-Coudert, Y. (2020) Oceanic thermal structure mediates dive sequences in a foraging seabird. Ecology and Evolution, 10, 6610–6622.

36. Oliver, E.C.J., Benthuysen, J.A., Bindoff, N.L., Hobday, A.J., Holbrook, N.J., Mundy, C.N. & Perkins-Kirkpatrick, S.E. (2017) The unprecedented 2015/16 Tasman Sea marine heatwave. Nature Communications, 8, 16101.

37. Ortega, J.C.G., Figueiredo, B.R.S., da Graça, W.J., Agostinho, A.A. & Bini, L.M. (2020) Negative effect of turbidity on prey capture for both visual and non-visual aquatic predators. The Journal of Animal Ecology, 89, 2427–2439.

38. Padleton, L.H., Aghmadia, G.N., Browman, H.I., Thurstand, R.H., Kaplan, D.M. & Bartolino, V. (2017) Debating the effectiveness of marine protected areas. ICES Journal of Marine Science.

39. Poloczanska, E.S., Babcock, R.C., Butler, A., Hobday, A., Hoegh-Guldberg, O., Kunz, T., Matear, R., Milton, D.A., Okey, T.A. & Richardson, A.J. (2007) Climate Change and Australian Marine Life. Oceanography and Marine Biology: An Annual Review,, 45, 407–478.

40. Pontzer, H. & McGrosky, A. (2022) Balancing growth, reproduction, maintenance, and activity in evolved energy economies. Current Biology, 32, R709–R719.

41. Pulvirenti, J., Reina, R.D. & Chiaradia, A. (2023) Exploring subcolony differences in foraging and reproductive success: the influence of environmental conditions on a central place foraging seabird. Royal Society Open Science, 10, 220362.

42. R Core Team. (2022) R: A Language and Environment for Statistical Computing.

43. Ramasco, V., Barraquand, F., Biuw, M., McConnell, B. & Nilssen, K.T. (2015) The intensity of horizontal and vertical search in a diving forager: the harbour seal. Movement Ecology, 3, 15.

44. Ramírez, F., Afán, I., Davis, L.S. & Chiaradia, A. (2017) Climate impacts on global hot spots of marine biodiversity. Science Advances, 3, e1601198.

45. Ramírez, F., Afán, I., Tavecchia, G., Catalán, I.A., Oro, D. & Sanz-Aguilar, A. (2016) Oceanographic drivers and mistiming processes shape breeding success in a seabird. Proceedings. Biological Sciences / the Royal Society, 283, 20152287.

46. Ropert-Coudert, Y., Chiaradia, A. & Kato, A. (2006) An exceptionally deep dive by a Little Penguin *Eudyptula minor*. Marine ornithology, 34, 71–74.

47. Ropert-Coudert, Y., Kato, A. & Chiaradia, A. (2009) Impact of small-scale environmental perturbations on local marine food resources: a case study of a predator, the little penguin. Proceedings. Biological Sciences / the Royal Society, 276, 4105–4109.

48. Ropert-Coudert, Y., Kato, A., Wilson, R.P. & Cannell, B. (2006) Foraging strategies and prey encounter rate of free-ranging Little Penguins. Marine Biology, 149, 139–148.

49. Sánchez, S., Reina, R.D., Kato, A., Ropert-Coudert, Y., Cavallo, C., Hays, G.C. & Chiaradia, A. (2018) Within-colony spatial segregation leads to foraging behaviour variation in a seabird. Marine Ecology Progress Series, 606, 215–230.

50. Saraux, C., Chiaradia, A., Salton, M., Dann, P. & Viblanc, V.A. (2016) Negative effects of wind speed on individual foraging performance and breeding success in little penguins. Ecological monographs, 86, 61–77.

51. Saraux, C., Robinson-Laverick, S.M., Le Maho, Y., Ropert-Coudert, Y. & Chiaradia, A. (2011) Plasticity in foraging strategies of inshore birds: how Little Penguins maintain body reserves while feeding offspring. Ecology, 92, 1909–1916.

52. Shin, Y.-J., Midgley, G.F., Archer, E.R.M., Arneth, A., Barnes, D.K.A., Chan, L., Hashimoto, S., Hoegh-Guldberg, O., Insarov, G., Leadley, P., Levin, L.A., Ngo, H.T., Pandit, R., Pires, A.P.F., Pörtner, H.-O., Rogers, A.D., Scholes, R.J., Settele, J. & Smith, P. (2022) Actions to halt biodiversity loss generally benefit the climate. Global Change Biology, 28, 2846–2874.

53. Soriano-Redondo, A., Inger, R., Sherley, R.B., Rees, E.C., Abadi, F., McElwaine, G., Colhoun, K., Einarsson, O., Thorstensen, S., Newth, J., Brides, K., Hodgson, D.J. & Bearhop, S. (2023) Demographic rates reveal the benefits of protected areas in a long-lived migratory bird. Proceedings of the National Academy of Sciences of the United States of America, 120, e2212035120.

54. Sutherland, D. & Dann, P. (2012) Improving the accuracy of population size estimates for burrow-nesting seabirds. Ibis, 154, 488–498.

55. Sutton, G.J. & Arnould, J.P.Y. (2022) Quantity over quality? Prey-field characteristics influence the foraging decisions of little penguins (Eudyptula minor). Royal Society Open Science, 9, 211171.

56. Wilson, R.P., Börger, L., Holton, M.D., Scantlebury, D.M., Gómez-Laich, A., Quintana, F., Rosell, F., Graf, P.M., Williams, H., Gunner, R., Hopkins, L., Marks, N., Geraldi, N.R., Duarte, C.M., Scott, R., Strano, M.S., Robotka, H., Eizaguirre, C., Fahlman, A. & Shepard, E.L.C. (2020) Estimates for energy expenditure in free-living animals using acceleration proxies: A reappraisal. The Journal of Animal Ecology, 89, 161– 172.

57. Wilson, R.P., Ropert-Coudert, Y. & Kato, A. (2002) Rush and grab strategies in foraging marine endotherms: the case for haste in penguins. Animal Behaviour, 63, 85–95.

58. Wood, S.N. (2011) Fast stable restricted maximum likelihood and marginal likelihood estimation of semiparametric generalized linear models. Journal of the Royal Statistical Society: Series B (Statistical Methodology), 73, 3–36.

59. Zentner, Y., Rovira, G., Margarit, N., Ortega, J., Casals, D., Medrano, A., Pagès-Escolà, M., Aspillaga, E., Capdevila, P., Figuerola-Ferrando, L., Riera, J.L., Hereu, B., Garrabou, J. & Linares, C. (2023) Marine protected areas in a changing ocean: Adaptive management can mitigate the synergistic effects of local and climate change impacts. Biological conservation, 282, 110048.

60. Zimmer, I., Ropert-Coudert, Y., Kato, A., Ancel, A. & Chiaradia, A. (2011) Does foraging performance change with age in female little penguins (Eudyptula minor)? Plos One, 6, e16098.

