## Supporting Information for "Decoding Ten Years of Little Penguin Foraging: Bio-Logging Reveals Foraging Patterns with Implications for Climate Change Mitigation and Marine Spatial Planning"

**Decadal-scale variation of little penguin foraging in the Bass Strait: insights from bio-loggers to assist Marine Spatial Planning**

Marianna Chimienti, Yan Ropert-Coudert, Benjamin Dupuis, Nicolas Joly, Claire Saraux, Andre Chiaradia, Akiko Kato

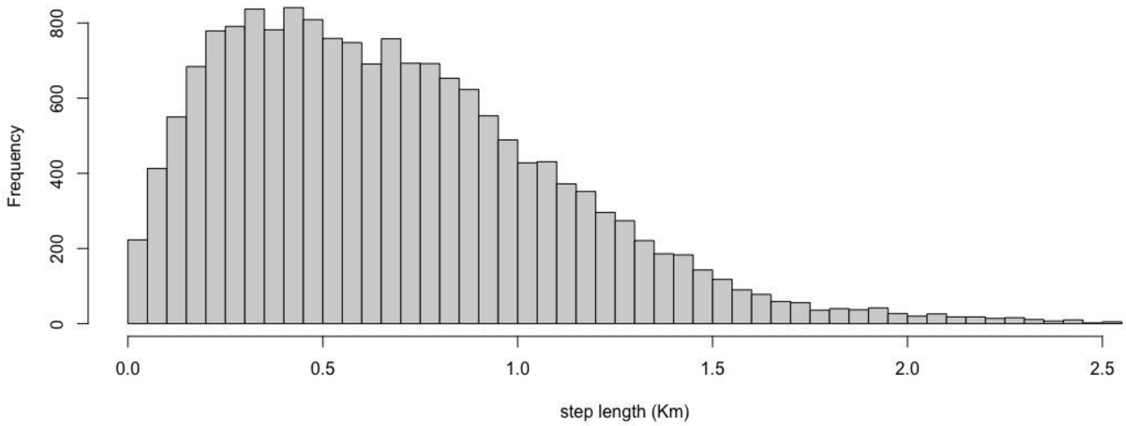


**Figure S1:** Distribution of step lengths performed by Little penguin (*Eudyptula minor*) obtained from the GPS dataset.

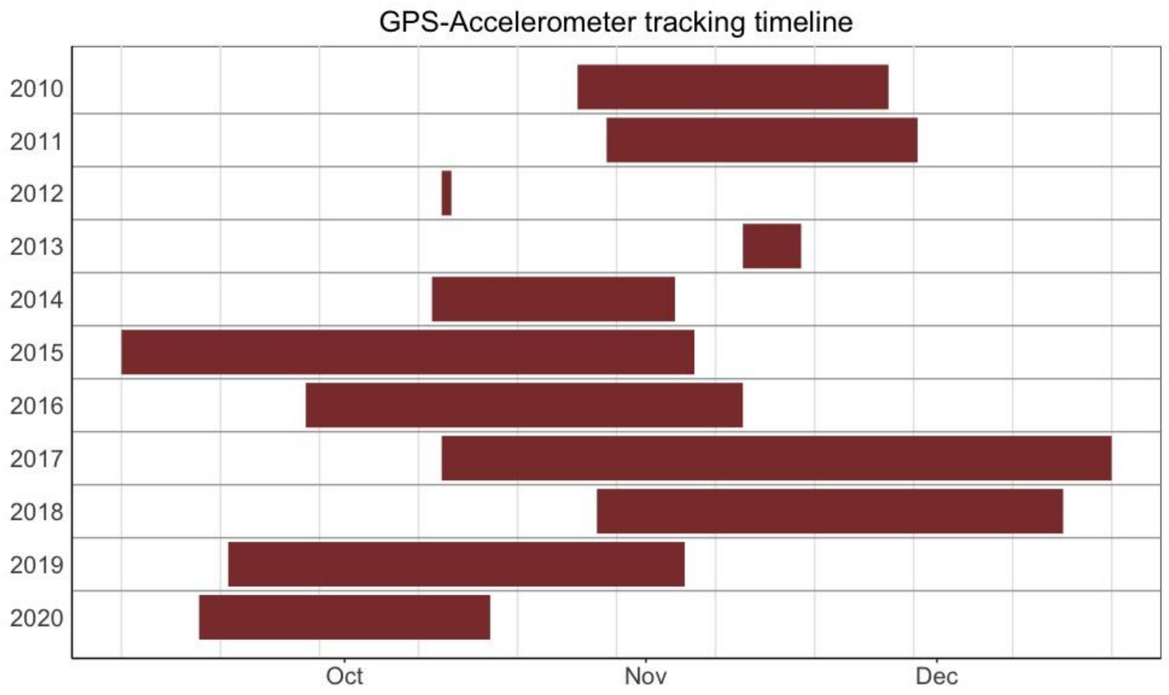


**Figure S2:** Time period of GPS-Accelerometer data availability for Little penguin (*Eudyptula minor*) across eleven sampling seasons.

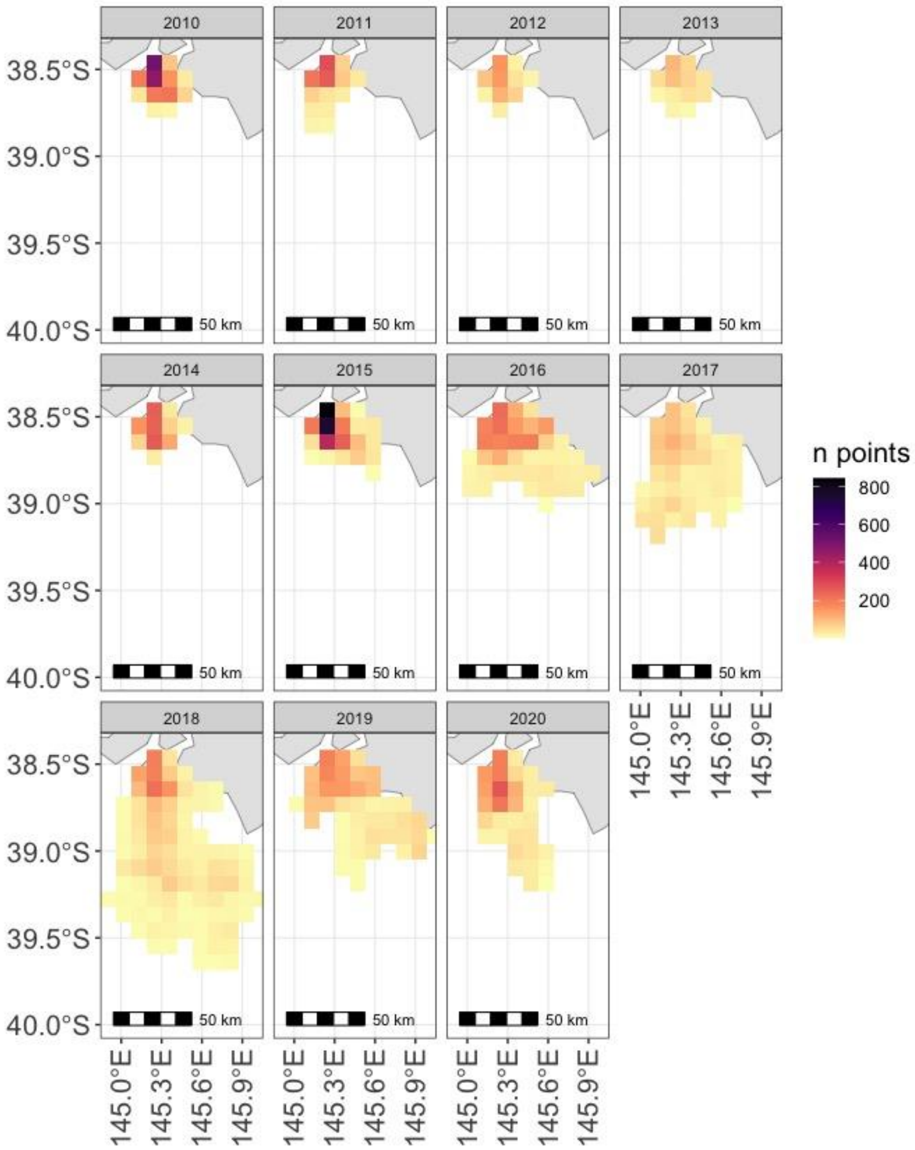


**Figure S3:** Yearly rasters summarizing the number of data points from GPS locations within each 10 Km$^2$ cell.

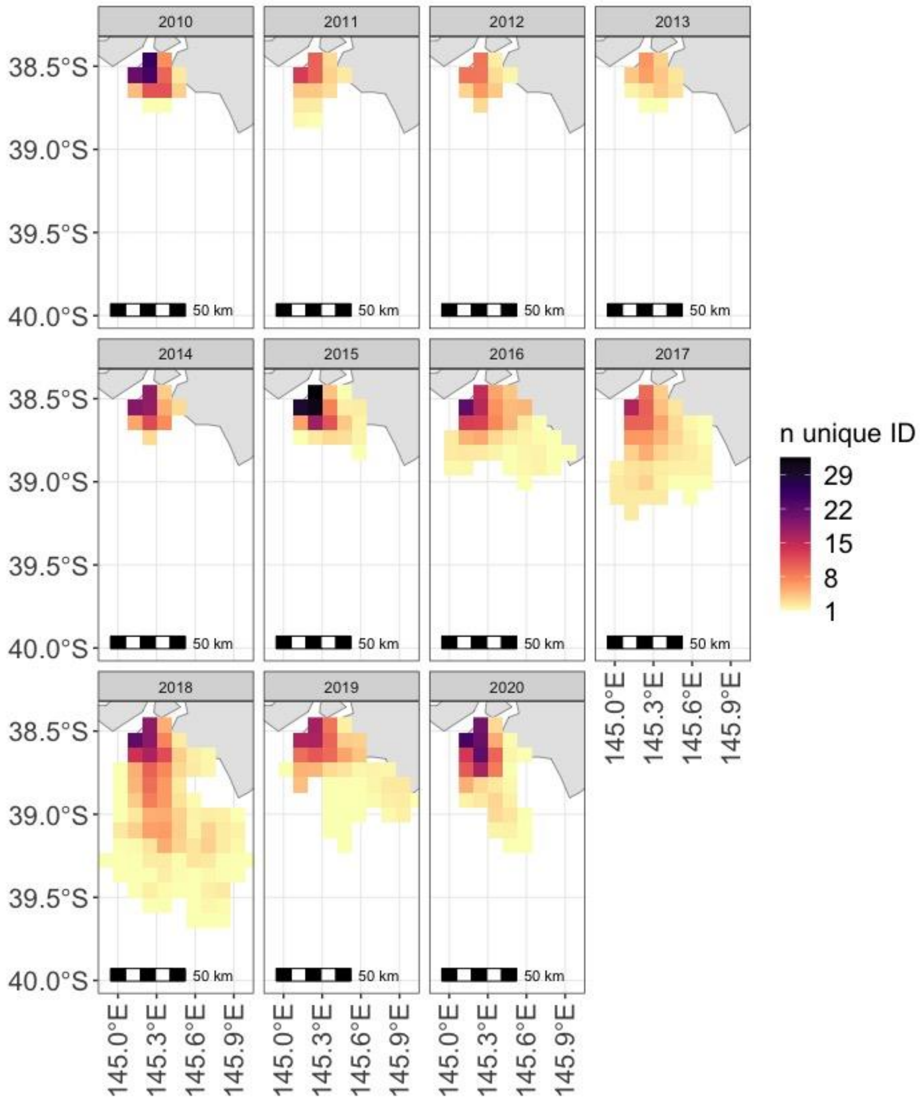


**Figure S4:** Yearly rasters summarizing the number of unique animals within each 10 $Km^2$ cell.

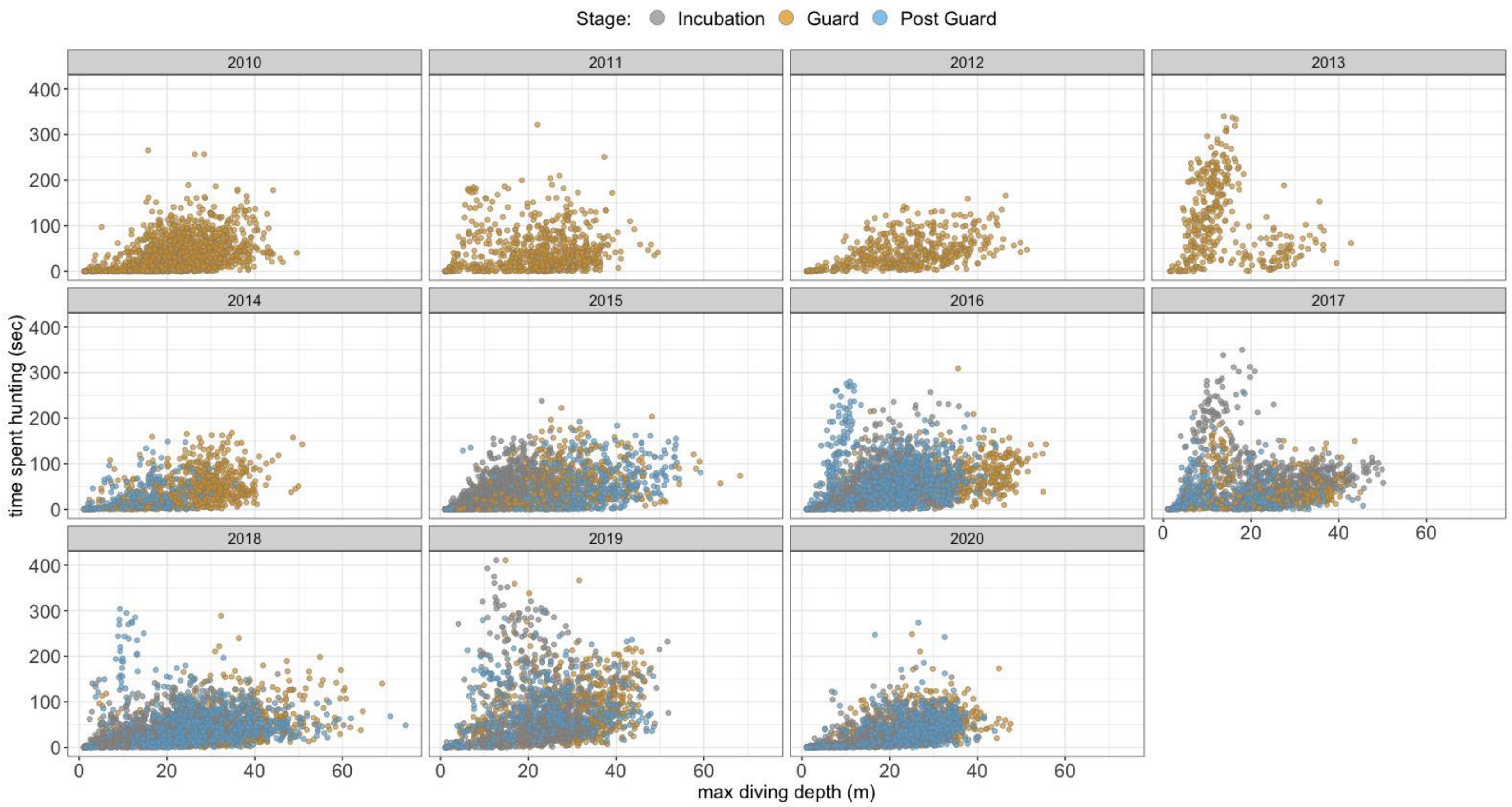


**Figure S5:** Time spent hunting (sec) versus maximum diving depth (m) across eleven breeding seasons. Colors indicate breeding stages: *Incubation* (grey), *Guard* (gold) and *Post Guard* (blue)

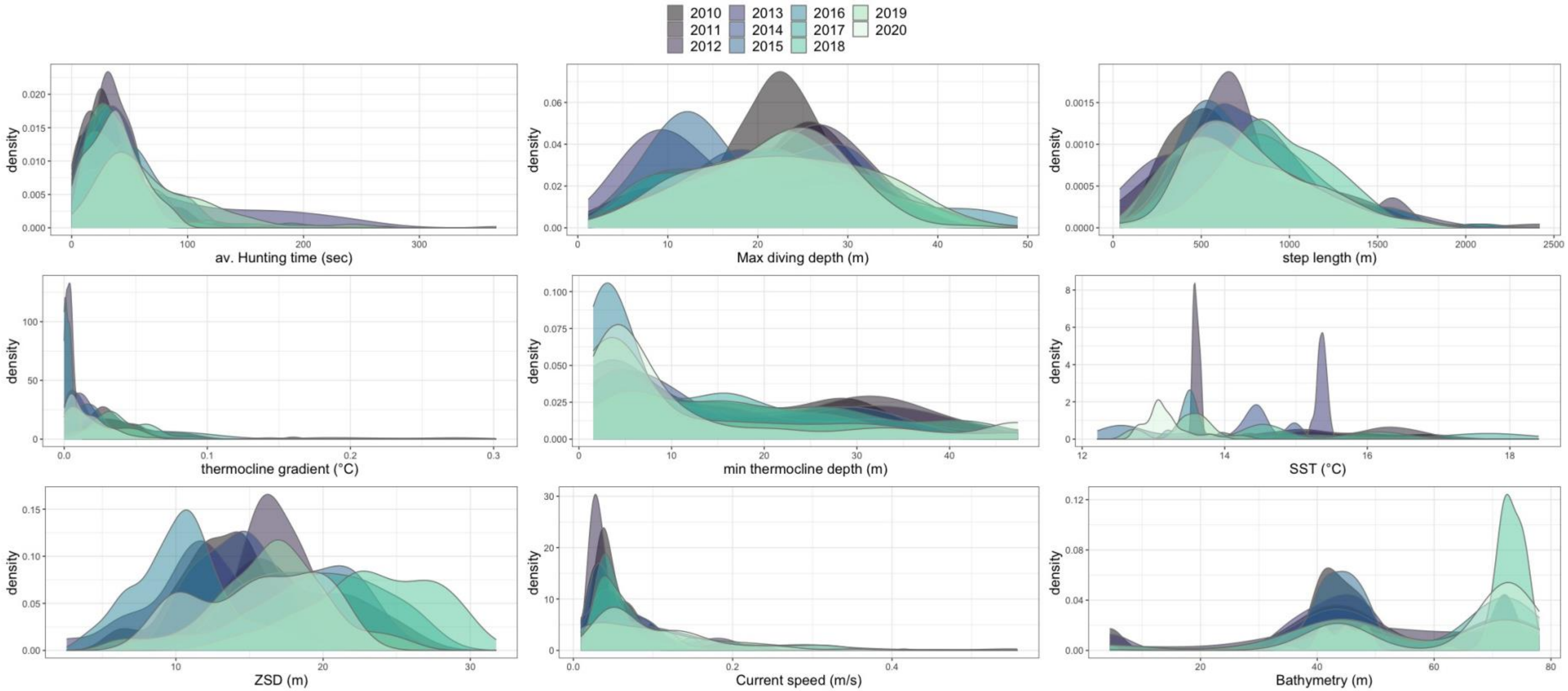


**Figure S6:** Yearly density distributions of environmental variables extracted from remote sensing datasets and matched with GPS-Accelerometer data collected on Little penguins breeding on Phillip Island (Australia).

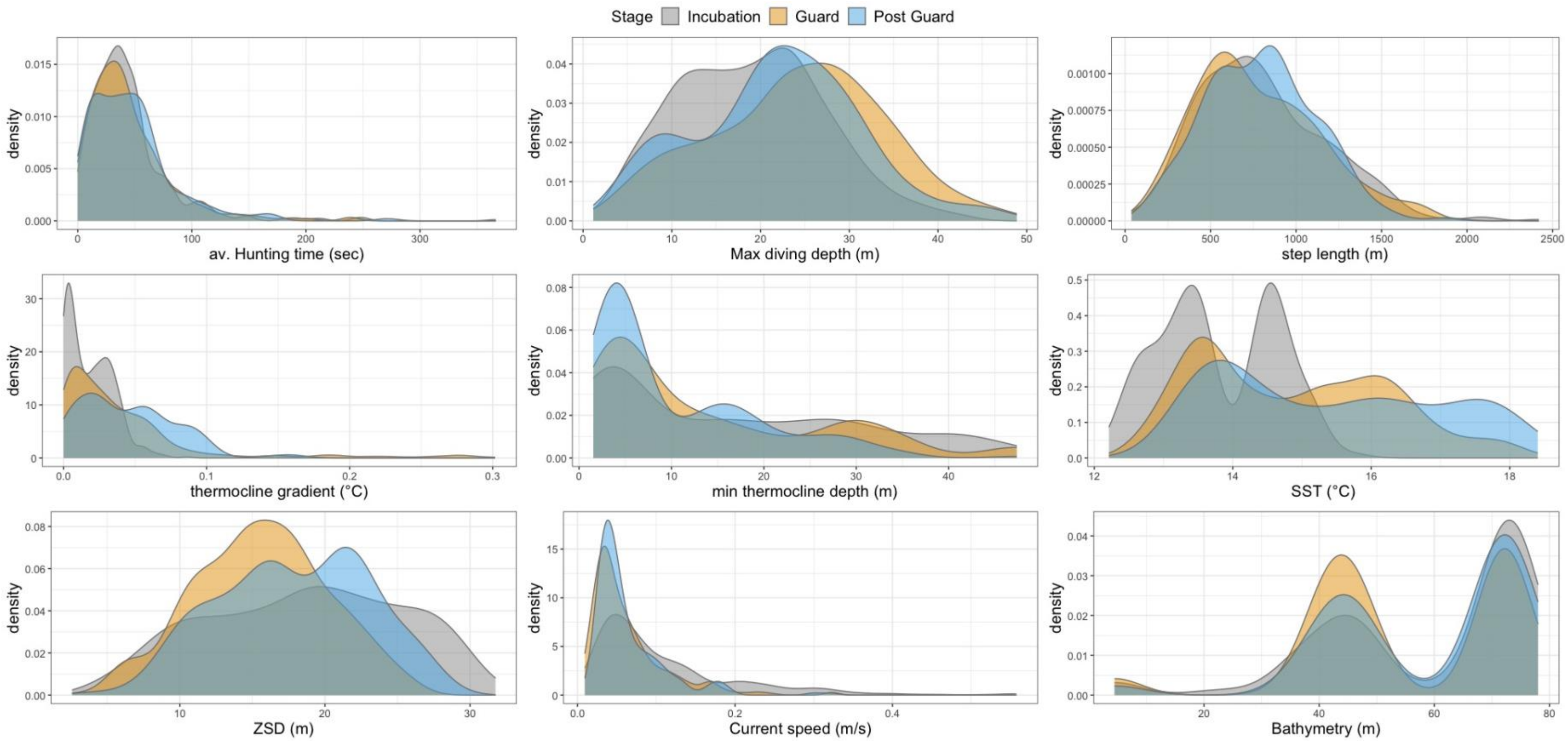


**Figure S7:** Density distributions of environmental variables extracted from remote sensing datasets and matched with GPS-Accelerometer data collected on Little penguins breeding on Phillip Island (Australia) grouped by breeding stage.

**Table S1:** Model selection showing the top five models. SST = Sea surface temperature, SZD = Secchi disk depth.

| **Formula** | **AICc** | **ΔAICc** |
|---|---|---|
| ~s(Max diving depth) + s(SST, by=Stage) + (step length)+ s(x,y) + s(SZD) + Stage + Season | 10495.9 | 0.00 |
| ~s(Max diving depth) + s(Bathymetry) + s(SST, by=Stage) + (step length) + s(x,y)+s(SZD) + Stage + Season | 10497.5 | 1.62 |
| ~s(Max diving depth) + s(SST, by=Stage) + (step length) + s(x,y) + Stage + Season | 10498.1 | 2.26 |
| ~s(Max diving depth) + s(SST, by=Stage) + (step length) + s(x,y)+s(SZD) + Season | 10498.3 | 2.45 |
| ~s(Max diving depth) + s(Bathymetry) + s(SST, by=Stage) + (step length) + s(x,y)+s(SZD) + Season | 10499.7 | 3.85 |

**Table S2:** Summary of the best model as selected in Table S1. Signif. codes: 0 '***' 0.001 '**' 0.01 '*' 0.05 '.' 0.1 ' ' 1

| **Parametric coefficients:** | | | |
|---|---|---|---|
| | **Estimate** | **Std. Error** | **p-value** |
| (Intercept-2010, Incubation) | 3.23502 | 0.12687 | *** |
| Stage - Guard | 0.05391 | 0.09646 | |
| Stage - Post Guard | 0.26230 | 0.11568 | * |
| Season 2011 | 0.50434 | 0.14506 | *** |
| Season 2012 | -0.14436 | 0.28952 | |
| Season 2013 | 1.04721 | 0.18456 | *** |
| Season 2014 | 0.01096 | 0.15177 | |
| Season 2015 | -0.02225 | 0.17195 | |
| Season 2016 | 0.15753 | 0.14465 | |
| Season 2017 | 0.73502 | 0.13633 | *** |
| Season 2018 | 0.63610 | 0.13517 | *** |
| Season 2019 | 0.62471 | 0.14651 | *** |
| Season 2020 | 0.05685 | 0.15291 | |

| **Approximate significance of smooth terms:** | | |
|---|---|---|
| | **edf** | **p-value** |
| s(max diving depth) | 8.138 | *** |
| s(SST): Incubation | 1.001 | |
| s(SST): Guard | 1.002 | *** |
| s(SST): Post Guard | 1.000 | *** |
| s(ZSD) | 1.901 | ** |
| s(step length) | 1.002 | *** |
| s(x,y) | 9.414 | *** |
